# A human *TSC1* mutation screening platform in GABAergic cortical interneurons for Genotype to Phenotype assessments

**DOI:** 10.1101/2020.06.01.128611

**Authors:** Dean Wundrach, Luis E Martinetti, April M Stafford, Stephanie M Bilinovich, Kartik Angara, Shane R Crandall, Daniel Vogt

## Abstract

Tuberous Sclerosis Complex is a complex syndrome that affects multiple organs and is caused by dysfunction of either the *TSC1* or *TSC2* genes. One of the least understood features of TSC is the impact of *TSC1&2* variants on brain phenotypes, including elevated rates of autism spectrum disorder and seizures. Moreover, while a great deal of work has uncovered how loss of either gene can alter various neural cell types, the impact of many variants in TSC and on these cell types is poorly understood. In particular, missense variants that cause minor changes in the proteins are expected to cause functional changes that differ from a complete loss of the protein. Herein, we examined how some missense variants in *TSC1* impacted the development of cortical inhibitory interneurons, a cell type whose molecular, cellular and physiological properties are known to be altered after loss of mouse *Tsc1*. Importantly, we found that most missense variants complemented phenotypes caused by loss of *Tsc1* and resulting in elevated MTOR activity as well as several cell intrinsic physiological properties. However, distinct variants showed deficits in complementing an increase in parvalbumin levels, which is observed after loss of *Tsc1* and demonstrated smaller amplitudes of after hyperpolarizations. These data suggest subtle but sensitive phenotypes can be detected by some *TSC1* missense variants and provide an *in vivo* system in which to better assess TSC variants.

## Introduction

The identification of genetic variants associated with neuropsychiatric disorders has elevated in recent years. This growth has been especially challenging in autism spectrum disorder (ASD), where several hundreds of genes have been implicated in the disorder, each with a plethora of different variants. The discovery of these genes involved in ASD has elevated our understanding of ASD molecular biology but opens the door for the need of novel assays to continue the advancement of knowledge (Rosti et al., 2014). While the functional impact of some variants can be easy to predict, i.e., loss of function, frameshift and nonsense, the impact of missense variants has been challenging to predict and validate.

The integration of more advanced variant support using computational tools to prioritize high impact variants with animal or humanized systems to define the physiological outcomes of the variant can build advanced genotype-to-phenotype insights. To validate these top variants an ideal method is to generate a knock-in animal model for each variant, which provides an *in vivo* environment for cells to develop. While ideal, this is costly, time consuming and inefficient, making it difficult to study variants in vivo. To more efficiently understand the impact of missense variants associated with ASD, we developed and validated an *in vivo* approach that can assess the impact of a variant in GABAergic cortical interneurons (CINs) (Vogt et al., 2015a, 2018).

CIN dysfunction is often implicated in ASD and CIN properties have been found to be altered in both humans diagnosed with ASD and in various ASD genetic deletion animal models (Pla et al.; Vogt et al., 2015a, 2018; Hoffman et al., 2016; Hashemi et al., 2017; Jung et al., 2017; Soghomonian et al., 2017; Elbert et al., 2019; Malik et al., 2019; Angara et al., 2020). CINs are derived from the medial and caudal ganglionic eminences (MGE and CGE), as well as the preoptic area (Wonders and Anderson, 2006; Gelman et al., 2011; Hu et al., 2017b). They tangentially migrate long distances to their final cortical destinations, laminate the cortex and begin to express unique molecular markers as they assume their diverse cell fates, each modulating cortical inhibition in exclusive ways (Wonders and Anderson, 2006; Miyoshi et al., 2010; Kessaris et al., 2014). The vast majority of CINs are derived from the MGE and can be delineated via the expression of somatostatin (SST) or parvalbumin (PV). Interestingly, PV expression is commonly affected in ASD and ASD gene animal models (Hashemi et al., 2017; Vogt et al., 2018; Malik et al., 2019), suggesting that this group of cells, and/or molecular marker, may provide a good readout of how ASD relevant missense variants could alter neural development.

Our previous work uncovered that genes underlying syndromes associated with high rates of ASD greatly impacted CIN development, and in turn, PV+ CINs (Vogt et al., 2014, 2015a, 2018; Pla et al., 2018), suggesting that these genes were essential for normal CIN development. Importantly, many of these genes regulate similar cellular processes, i.e., mammalian target of rapamycin (MTOR) signaling. One of these genes, *TSC1*, inhibits MTOR activity and, when mutated, results in the syndrome, Tuberous Sclerosis Complex (TSC). Notably, conditional loss of *Tsc1* in mouse CINs leads to ectopic expression of PV and fast-spiking physiological properties (Malik et al., 2019). However, nothing is known about how the multitude of variants in *TSC1* could impact CIN development and the molecular and physiological properties of PV+ CINs. We developed a platform to test genomic human variants of *TSC1* within these CINs using a cell specific knockout followed by human allele recovery. A selection of *TSC1* variants that are low allele frequency throughout the population (based on gnomAD) were tested to study their involvement in altering cell function. The variants chosen have conflicting reports for their involvement in ASD, where early studies identified the variants in patients with ASD (Schaaf et al., 2011; Kelleher et al., 2012), while more recent data suggests either no disease association or a more complex multifactorial role in disease, making them great candidates for our *in vivo* assay. While all variants were able to complement classical MTOR activity phenotypes, distinct variants failed to complement the elevated PV expression associated with loss of *Tsc1*. While many physiological properties were unchanged by one of these variants, we did uncover decreased action potential after hyperpolarizations, suggesting subtle phenotypes that may associate with multifactorial ASD development. These data demonstrate a sensitive readout for TSC variant function *in vivo* and suggest the further expansion of this model into more challenging TSC variants, such as variants of unknown significance (VUS).

## Materials and methods

### Animals

*Tsc1^Flox^* (Kwiatkowski et al., 2002) and *Ai14* (Madisen et al., 2010) have been previously described. Both lines were back crossed to a CD-1 background for at least five generations before experiments began. For timed matings, noon on the day of the vaginal plug was considered embryonic day 0.5. Experimenters were blind to the genotypes of the mice and littermates were used as controls when possible. Since our previous work did not find a difference in sex phenotypes (Malik et al., 2019), both sexes were used. All mouse procedures were performed in accordance with NIH Guidelines for the Care and Use of Laboratory Animals and were approved by the Michigan State University Institutional Animal Care and Use Committee.

### DNA vector generation

The *DlxI12b-BG-hTSC1-IRES-Cre* lentiviral DNA vector was previously described (Malik et al., 2019). To generate the ASD-relevant *hTSC1* variants, we designed gene blocks (integrated DNA technologies) that included each mutation and flanking endogenous restriction sites that resided within the human *TSC1* gene. Next, the gene blocks were ligated into the *DlxI12b-BG-hTSC1-IRES-Cre* vector (replacing the WT sequence), and then verified using Sanger sequencing.

### *In vitro* slice preparation

Coronal cortical slices (300 μm thick) were prepared from mice (between postnatal ages 34 and 56) using methods previously described (Crandall et al., 2015, 2017). Briefly, mice were deeply anesthetized with isoflurane before decapitation. Brains were then quickly removed and placed in a cold (~4°C) oxygenated slicing solution (95% O2, 5% CO2) containing (in mM): 3 KCl, 1.25 NaH2PO4, 10 MgSO4, 0.5 CaCl2, 26 NaHCO3, 10 glucose and 234 sucrose. Slices were cut using a vibrating tissue microtome (Leica VT1200S) and then transferred into a holding chamber containing warm (32°C) oxygenated (95% O2, 5% CO2) artificial cerebrospinal fluid (ACSF) containing (in mM): 126 NaCl, 3 KCl, 1.25 NaH2PO4, 2 MgSO4, 2 CaCl2, 26 NaHCO3 and 10 glucose. Slices were kept at 32°C for 20 min followed by room temperature for an additional 40 min before recording.

### *In vitro* electrophysiological recordings, data acquisition, and analysis

For recordings, individual slices were transferred to a submersion recording chamber and continually perfused (~3 ml/min) with warm (32°C) oxygenated (95% O2, 5% CO2) ACSF containing (in mM): 126 NaCl, 3 KCl, 1.25 NaH2PO4, 2 MgSO4, 2 CaCl2, 26 NaHCO3 and 10 glucose. Neurons were visualized using infrared differential interference contrast (IR-DIC) and fluorescence imaging using a Zeiss Axio Examiner.A1 microscope mounted with a video camera (Olympus XM10-IR) and a 40x water-immersion objective. Cells expressing tdTomato were randomly targeted in cortex for patching. Whole-cell recordings were obtained using borosilicate glass pipettes (4-6 MΩ tip resistance) containing a potassium-based internal solution (in mM): 130 K-gluconate, 4 KCl, 2NaCl, 10 HEPES, 0.2 EGTA, 4 ATP-Mg, 0.3 GTP-Tris and 14 phosphocreatine-K (pH 7.25, 290 mOsm). All whole-cell recordings were corrected for a 14-mV liquid junction potential.

Electrophysiological data were recorded and digitized at 20 or 50 kHz using Molecular Devices hardware and software (MultiClamp 700B amplifier, Digidata 1550B4, and pClamp11.1). Signals were low-pass filtered at 10 kHz prior to digitizing. During the recordings, the pipette capacitances were neutralized and series resistances (typically between 10-25 MΩ) were compensated online (100% for current-clamp). The series resistances were continually monitored throughout the recordings.

Analysis of *in vitro* electrophysiological data was performed in Molecular Devices and Microsoft Excel using previously described methods (Crandall et al., 2017). Briefly, resting membrane potentials (RMP, in mV) were measure immediately after break-in, with no applied current. Input resistance (Rin, in MΩ) was measured using Ohm’s law by measuring the voltage response from rest to an injection of a small negative current (5-20 pA). Membrane time constants (τm, in ms) were measured from the average response (see Rin above) by fitting a single exponential to the initial falling phase of the response (100-300 ms; omitting the 1st ms). Membrane capacitance (Cin, in pF) was calculated by τm / Rin. Rheobase currents (in pA) were defined as the minimum positive current (5 pA steps) to elicit an action potential (AP) from a holding potential of −79 mV. APs properties were measured from the first spike evoked by the rheobase current. AP threshold (in mV) was defined as the membrane potential at which its first derivitive (*d*V/*d*t) exceeded 10 mV/ms. AP amplitudes (in mV) were defined as the voltage difference between the threshold and the peak of the AP. AP half-widths (in ms) were measured at the half-height between the threshold and the AP peak. The max rate of rise (in mV/ms) was defined as the maximal *d*V/*d*t during the rising phase of the AP, whereas the max rate of decay (in mV/ms) was defined as the maximal negative *d*V/*d*t during the falling phase of the same AP. Fast afterhyperpolarization potentials (fAHPs, in mV) were measured as the difference between the AP threshold and the peak negative potential of the AHP immediately following the AP. Post train medium afterhyperpolarization potentials (mAHPs, in mV) were measured as the difference between a baseline period (500 ms) prior to current injection and the peak negative potential following the 1 sec train of APs. Analyses were performed on the first suprathreshold current injection in which the initial firing frequency of the cell exceeded 150 Hz, and at least 100 APs were evoked. Membrane potential sags (in mV) were measured using a 1 s negative current step that hyperpolarized the neuron from −79 to −99 mV and were calculated relative to the steady-state voltage at the end of the step. Frequency-intensity (F/I) relationships were obtained by holding the soma at −79 mV with intracellular current and injecting suprathreshold positive current (50 pA steps, 1 s duration). F/I slopes (in Hz/pA) were determined using the initial frequency (reciprocal of the first interspike interval) over the entire F-I plot. Spike frequency adaptation was determined by calculating the adaptation ratio, defined as the steady-state firing frequency (average of the last 5 APs) divided by the initial frequency. Spike height accommodation was defined as the amplitude of the last AP divided by the first AP. Analysis for both the spike frequency adaptation and spike height accommodation was performed on the first suprathreshold current injection in which the initial firing frequency exceeded 150 Hz.

### Immuno-fluorescence labeling and imaging

Primary neurons and coronal brain sections were washed in PBS containing 0.3% Triton-X100, blocked in the same solution containing 5% BSA and then incubated in primary antibodies for 1-2 hours (or overnight). They were then washed 3 times and then incubated with secondary antibodies containing fluorophores for 1 hour before 3 final washes. Primary neurons were stored in PBS while sections were cover slipped. Primary antibodies included rabbit anti-GABA 1:500 (Sigma A2052), rabbit anti-parvalbumin 1:400 (Swant, PV-27). Alexa-conjugated fluorescent secondary antibodies (Thermo-Fisher) were used to detect reactivity of primary antibodies. Native tdTomato fluorescence was imaged and in vitro primary MGE cultures were also labeled for DAPI using NucBlue Fixed Cell ReadyProbes (Thermo Fisher, R37609). MGE primary cultures were imaged using a Nikon eclipse Ts2R microscope with a Photometrics coolsnap dyno camera. Transplant tissue sections were imaged using a Leica DM2000 compound microscope with an attached camera (DFC3000G).

### Lentivirus preparation

Lentiviruses were prepared as previously described (Vogt et al., 2015b). Briefly, the *TSC1* lentiviral vectors were co-transfected with *pVSV-g*, *pRSVr* and *pMDLg-pRRE* helper plasmids using Lipofectamine^2000^ (Thermo Fisher Scientific) into HEK293T cells, and the media replaced after four hours. On the fourth day after transfection, media was collected and filtered to remove cells and debris and then complexed with Lenti-X concentrator (Clontech) according to the manufacturer’s protocol to concentrate lentiviral particles. Concentrated lentiviruses were stored at −80°C until use.

### MGE primary cultures

We performed MGE primary cultures as described in (Angara et al., 2020). Briefly, we cultured the MGE cells in DMEM supplemented with 10% FBS and penicillin/streptomycin from time of seeding until one day *in vitro*. The cells were transduced with virus at this stage for four hours. After four hours of transduction, we replaced all media (and removed any excess virus) with neurobasal media, supplemented with B27, glucose, glutamax and penicillin/streptomycin, and left the cells in this media until seven days *in vitro*. At that time, the cells were fixed in 4% paraformaldehyde and then assessed via immuno-fluorescence.

### MGE transplantation

MGE cells that were transduced with lentivirus and transplanted into neonatal mouse cortices were performed as previously described (Vogt et al., 2015b). Briefly, *Ai14^Flox/+^* E13.5 MGE cells that were either WT, *Tsc1^Flox/+^* or *Tsc1^Flox/Flox^* were transduced with *hTSC1-Cre* lentiviruses, and then transplanted into WT mouse neonatal cortices at multiple sites. The cells developed *in vivo* until 35 days post-transplant. After this time, cells were identified by native tdTomato expression and were co-labeled for molecular markers via immuno-fluorescence.

### Statistics and cell assessments

Graphpad Prism 7 and Origin Pro 2019 were used to calculate statistical significance; a p value of <0.05 was considered significant. For data with parametric measurements, we used a One-Way ANOVA with a Tukey post-test or a Two-sample t test to determine significance. For non-parametric data sets (transplantation experiments where data were normalized), we used a Chi-squared test to determine significance or a Mann–Whitney U test.

### Western blotting

Cell pellets were lysed using RIPA buffer. Each sample lysate was diluted to obtain a concentration of 1mg/mL in a buffer containg protein loading dye. 10uL of the protein lysate was loaded into a bis-tris protein gel (4-12% Invitrogen Bolt Bis-Tris mini gel), ran at 150V for 35min then transferred to PVDF membrane using the iBlot2 system. Blots were blocked with 5% milk in TBST for 1 hour prior to incubation with primary antibody (1:1000 in TBST + 5% BSA) for 1 hour at room temperature. Blots were washed 4 times with TBST and incubated with secondary antibody (1:2000, HRP-conjugated anti rabbit, BioRad) for 1hr at room temperature then washed 4 times with TBST. Signal was detected using SuperSignal West Femto (ThermoScientific) chemiluminescent substrate and imaged on a ChemiDoc system (BioRad). Primary antibodies included rabbit anti-HAMARTIN (Cell Signaling Technologies, 4906) and rabbit anti-GAPDH (Cell Signaling Technologies, 2118).

## Results

### Generation and validation of human *TSC1* variants associated with ASD

We chose five human (h)*TSC1* variants based on their past suggested association with an ASD co-diagnosis (Schaaf et al., 2011; Kelleher et al., 2012); R336W, T360N, T393I, S403L and H732Y, and subcloned into a lentiviral DNA vector (Figure 1A, top). All five variants are annotated as either conflicting or benign in ClinVar. Thus, these variants are especially difficult to understand whether they will impact cellular phenotypes and require analyses with our *in vivo* complementation assay.

**Figure 1:**
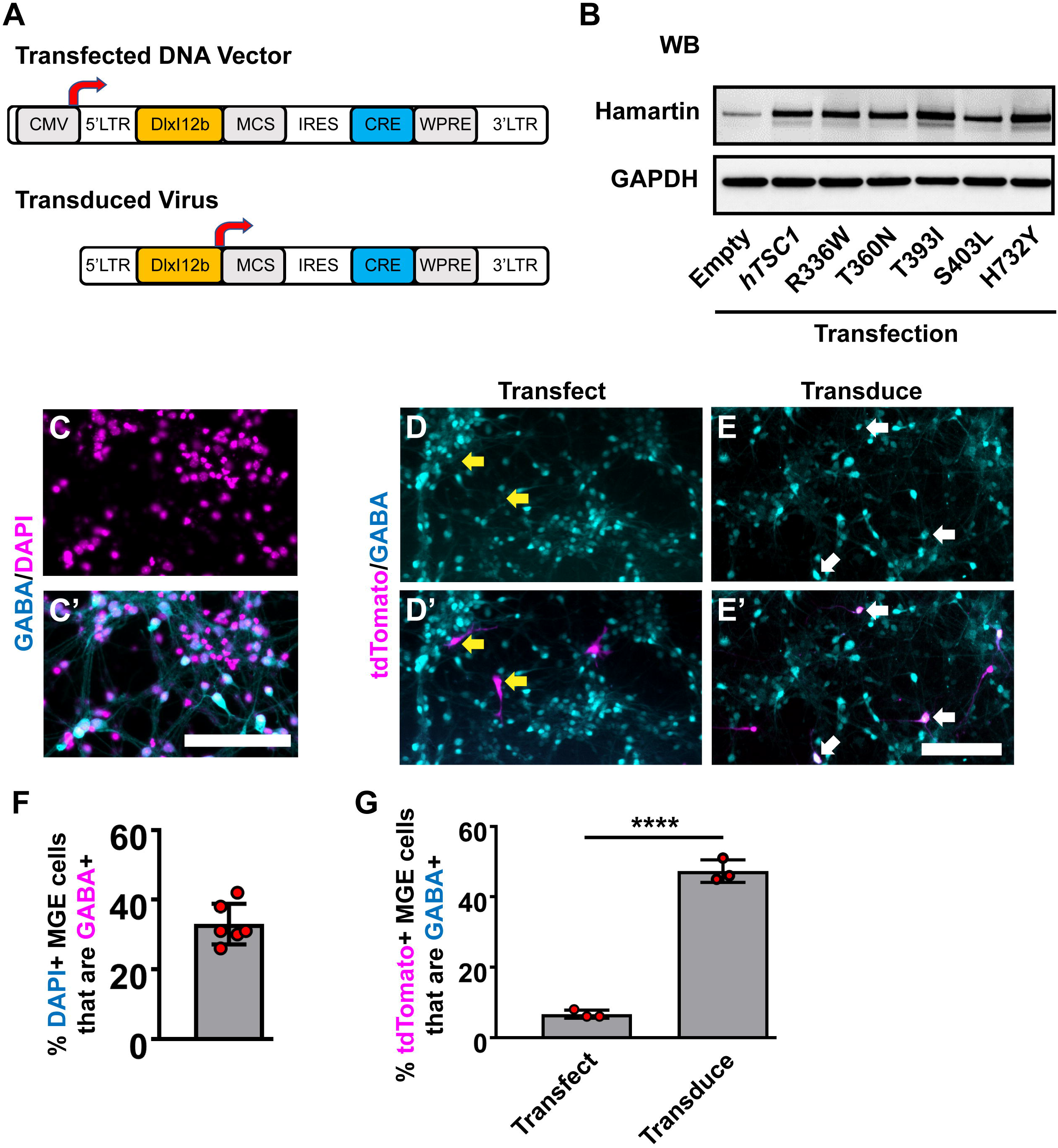
Human *TSC1* variants and expression in MGE primary cultures. (A) Schema depicting the lentiviral DNA vector and resulting lentivirus to express human *TSC1* variants and Cre recombinase. (B) *TSC1* variants expressed in HEK293T cells show increased levels of expression over endogenous HAMARTIN protein and migration at the correct molecular weight. (C, C’) E13.5 MGE primary cultures that were grown for 7 days and co-labeled for GABA and DAPI. E13.5 MGE primary cultures that were either transfected (D, D’) or transduced (E, E’) with the vector or virus depicted in (A) and co-labeled for the Cre-dependent reporter, tdTomato, and GABA. (F) Quantification of the proportion of DAPI+ cells that express GABA in MGE cultures. (G) Quantification of the proportion of tdTomato+ cells that are GABA+. Yellow arrows denote non co-labeled cells while white arrows denote co-labeled cells. Data are expressed as the mean ± SEM. Data were collected from 3 biological replicates (transplants) for all groups. For soma size a total of 75 cells were counted for each group and for PV labeling the number of tdTomato+ CINs assessed were: Cre only 454, WT 152, R336W 205, T360N 156, T393I 295, S403L 148 and H732Y 137. Abbreviations: (WB) western blot; (KD) kilodalton. Scale bars in (C’ and E’) = 100 μm.

To this end, we first assessed the expression of these variants in our lentiviral vector. While the DNA vector has the ability to drive expression from a strong CMV promoter, the resulting virus uses the *DlxI12b* enhancer, which biases expression to GABAergic neurons (Arguello et al., 2013). Each version expresses either Cre recombinase alone or in combination with WT *hTSC1* or each of the variants. To assess if each of the variants generated proteins at the correct molecular weight, we transfected each into HEK293T cells and assessed for the *TSC1* encoded protein, Hamartin. Both the WT and variants expressed at elevated levels over the endogenous protein and at the correct size, indicating that we could express both WT and variants (Figure 1B).

Next, we utilized MGE primary cultures to test these vectors and viruses *in vitro*. Since cells in the developing MGE include both GABAergic neurons and other types of cells, we asked what proportion of cells were GABAergic in primary E13.5 MGE cultures that had grown *in vitro* for 7 days. Roughly 30-40% of the cells were GABAergic (Figures 1C, 1C’). We either transfected the WT *hTSC1* expression vector or transduced the virus into the MGE cultures 24 hours after seeding and then assessed how many GABAergic cells co-labeled with tdTomato (Cre-expressing) after 6 additional days. Only ~6% of transfected cells co-labeled for GABA and tdTomato, despite 30-40% of the cultures being GABA+ (Figures 1D, 1D’, 1G). However, the *DlxI12b* virus transduced cells had ~50% co-labeled cells. Thus, utilizing the *DlxI12b* enhancer virus was more efficient at biasing towards GABAergic cells but can not override the strong CMV promoter in the DNA vector. Finally, we transduced the various viruses into the MGE cultures in the same manner but by day 7 *in vitro* did not observe gross differences in cell morphology (data not shown), suggesting that a longer developmental timeline was warranted.

### *In vivo* assay to determine the impact of *hTSC1* missense variants

To fully understand the impact of *hTSC1* variants in MGE-lineage CIN development and function, we modified a validated *in vivo* assay (Vogt et al., 2015b) to assess each variant individually. This assay also allows for MGE cells to develop over a long period after being genetically manipulated by the viruses. To this end, we transduced E13.5 MGE cells that were *Tsc1^Flox/Flox^*; *Ai14^Flox/+^* with a lentivirus carrying genes for *Cre* and either nothing (Empty), a WT *hTSC1* or each of the missense variants (Schema Figure 2A). After transduction, the cells (now tdTomato+) were transplanted into a postnatal day (P)1 WT pup to develop *in vivo*. TdTomato+ transplants were then assessed 35 days later for soma size and PV expression, which are both increased after deletion of mouse *Tsc1* in CINs (Malik et al., 2019). Consistent with our previous findings, using a Cre only virus to delete mouse *Tsc1* from MGE-lineage CINs, resulted in elevated soma sizes and over 40% of CINs expressing PV (Figures. 2B, 2I and 2J). Expression of WT *hTSC1* complemented these phenotypes, i.e. reduced soma size and decreased PV expression (Figures. 2C, 2I and 2J). Notably, all five variants complemented the increase in soma size induced by loss of *Tsc1* (vs. Cre, p<0.0001 for all variants except H732>Y, which was p = 0.0001), and all variants resembled WT levels (Figures. 2B–2I).

**Figure 2:**
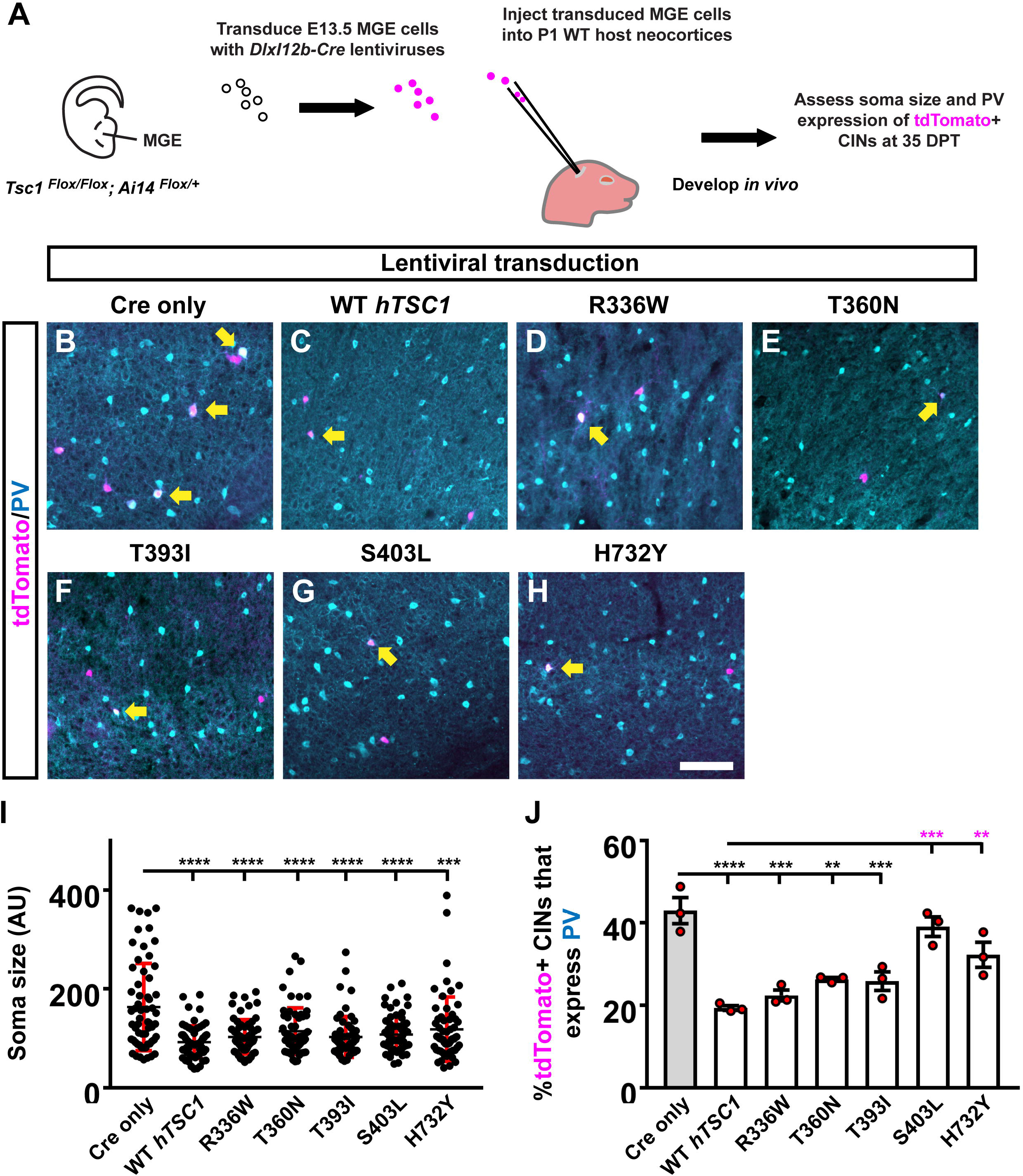
Distinct *TSC1* variants impact PV expression but not soma size. Schema depicting the complementation assay (A). Briefly, *Tsc1^Flox/Flox^; Ai14^Flox/Flox^* E13.5 MGE cells were dissociated, transduced with Cre-expressing viruses, transplanted into WT neocortices and assessed after developing *in vivo* for 35 days. (B-H) Example immuno-fluorescent images of transplanted CINs in the neocortex co-labeled for tdTomato and PV. Arrows point to co-labeled cells. (I) Quantification of the soma size for each variant complementation. (AU) arbitrary units. (J) Quantification of the %tdTomato+ CINs that express PV. Data are expressed as the mean ± SD for soma size and SEM for PV counts, n = 3 for all groups. **p<0.01; ***p<0.001; ****p<0.0001. Scale bar in (H) = 100 μm.

While increased soma size is a known phenotype associated with the loss of *Tsc* gene function, it often only occurs with the complete ablation of both *Tsc* genes (Tavazoie et al., 2005; Malik et al., 2019), suggesting that these missense variants are functional. Since we found some intermediate phenotypes in PV expression in our *Tsc1* heterozygous studies (Malik et al., 2019), we asked whether expression of PV may be a more sensitive readout to screen the impact of *hTSC1* missense variants. In our complementation assay, we found that PV expression and physiological properties associated with PV-expressing CINs are often altered in other syndromic ASD animal models and by missense mutations in other key ASD risk genes (Vogt et al., 2015a, 2018). Thus, we assessed the expression of PV in the transduced MGE cells. Of the five missense variants, there were two that did not complement the increase in PV expression, i.e. S403L and H732Y (Figs. 2G, 2H and 2J; S403L, p = 0.0003, H732Y, p = 0.004), suggesting that these variants partially impact CIN molecular properties while still being able to complement the increase in cell size. While the other variants were not significantly different than WT *hTSC1*, they, like the WT version, were significantly different than Cre-mediated loss of mouse *Tsc1* (Figs. 2B–2F and 2J; WT, p < 0.0001, R336W, p = 0.0002, T360N, p = 0.009, T393I, p = 0.0007). Overall, these data demonstrate that PV expression may be a sensitive TSC phenotype and that distinct variants are more likely to affect CIN molecular properties.

### Afterhyperpolariztions are abnormal in S403L CINs with FS properties

The greater propensity of CINs with the S403L mutation to express PV when compared to WT *hTSC1* could have an impact on the electrophysiological properties of these neurons. PV is an EF-hand calcium-binding protein that is found in distinct classes of neurons in the mammalian neocortex and is likely to play an essential physiological function in the behavior of these cells (Celio, 1986). To investigate the functional effects of the S403L variant, we made whole-cell recordings of transduced MGE CINs in the somatosensory cortex from acute coronal slices obtained from similarly aged littermate mice transplanted with cells carrying genes for either an S403L missense variant or the WT *hTSC1* protein. Transduced cells were identified by their tdTomato expression (Figure 3A). As a population, the passive and active membrane properties of S403L CINs were not significantly different from WT *hTSC1* cells (Supplemental Table 1).

**Figure 3:**
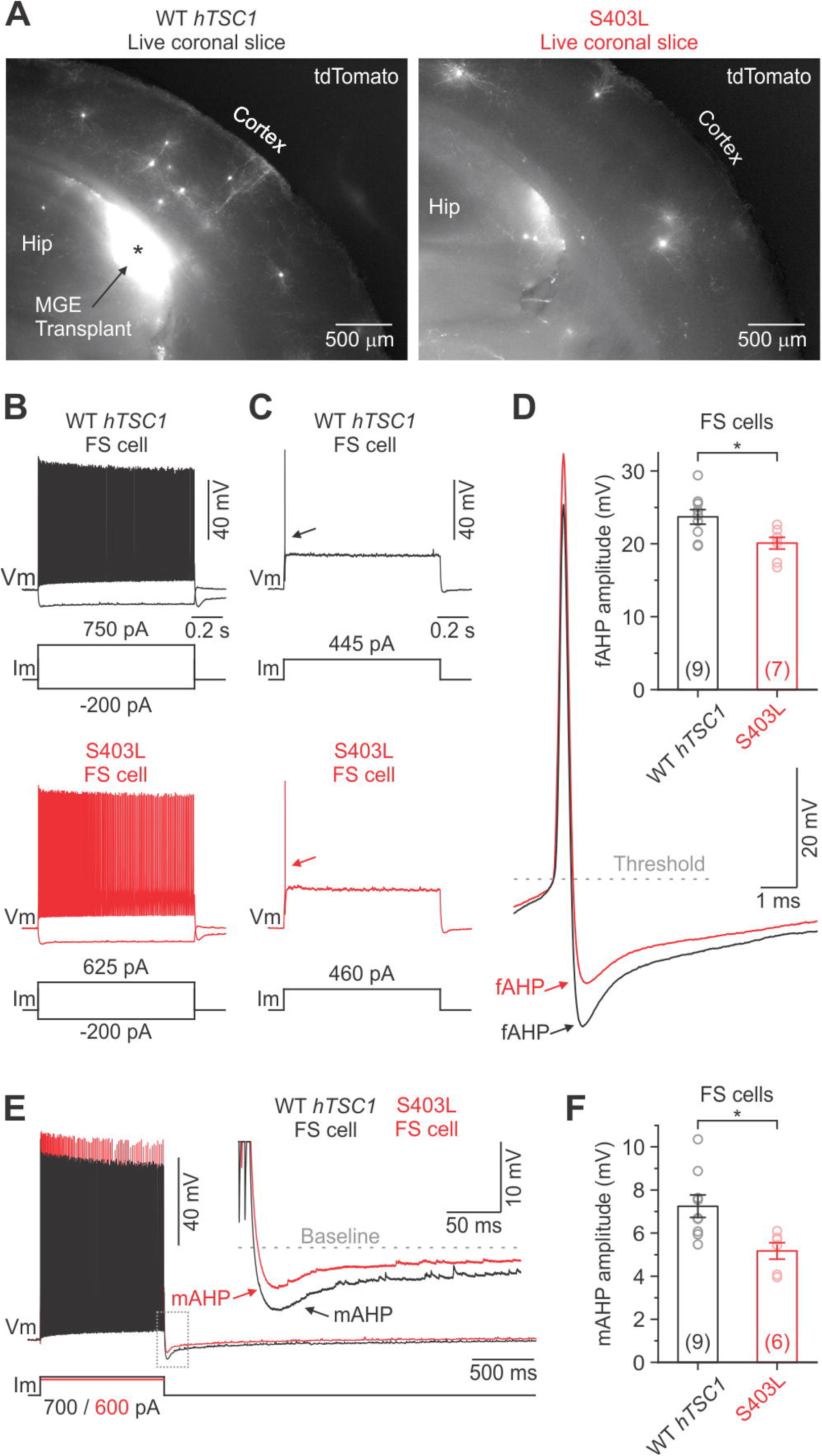
Fast and medium afterhyperpolarizations are reduced in S403L CINs with FS properties. (A) Low-magnification fluorescence images taken of live coronal brain sections (300 μm) showing tdTomato-expressing MGE cells transduced with either the WT *TSC1* or S403L missense variant after developing *in vivo*. (B) Voltage responses of a WT *TSC1* and S403L CIN to intracellular current steps (1 sec). Both cells were classified as FS using a non-linear classifier based on key electrophysiological properties (see results). (C) Responses to a threshold current injection (rheobase) for the same cells shown in (B). (D) Left, superimposed are the indicated single APs from (C: arrows) at increased magnification. Note the smaller fAHP amplitude immediately following the AP for the S403L cell. fAHPs were measured as the difference between the AP threshold and the peak negative potential. Right, quantification of the fAHP amplitude for each variant (fAHP WT *TSC1*: 23.7 ± 1.0 ms, n = 9 cells from 4 mice; fAHP S403L: 20.1 ± 0.8 ms, n = 7 cells from 4 mice; p < 0.02, two-sample t-test, two-tailed). (E) Spike trains evoked by a positive injected current. Traces are from the same cells shown in (B). Inset, showing the post-train mAHP at increased magnification. Note the smaller mAHP amplitude immediately following the spike train for the S403>L cell. (F) Quantification of the post-train mAHP amplitude for each variant. (mAHP WT *TSC1*: 7.2 ± 0.5 ms, n = 9 cells from 4 mice; mAHP S403L: 5.2 ± 0.4 ms, n = 6 cells from 4 mice; p < 0.02, two-sample t-test, two-tailed). Data are expressed as the mean ± SEM. *p<0.05.

Since deletion of mouse *Tsc1* can aberrantly lead to FS properties (Malik et al., 2019), we asked if the S403L variant led to more CINs with FS properties. To verify the cell identity objectively, we used a simple non-linear classifier based on three electrophysiological properties, as previously described (Hu et al., 2013). These parameters consisted of action potential half-width (APHW), afterhyperpolarization amplitude (AHP), and spike frequency adaptation ratio (AR). These properties are typical of FS CINs and distinct from that of SST expressing CINs, the other major class of MGE-derived CINs (Wonders and Anderson, 2006). Cells were classified as FS if at least two of the following conditions were true: APHW < 0.26 ms, AHP > 16.5 mV, and AR > 0.56. If cells did not meet these conditions, they were classified as non-FS cells (i.e., putative SST-expressing CINs). When we applied the classifier to our population of S403L CINs, we found that 56.3 ± 9.2 % of the cells were classified as FS (n = 4 mice). Data from WT *hTSC1* CINs, however, yielded a similar percentage of FS cells (45.0 ± 12.6 %; n = 5 mice; p = 0.52, two-sample t-test, two-tailed).

In regard to intrinsic membrane properties, S403L FS cells did not differ from WT *hTSC1* FS cells in either their resting potential, input resistance, membrane time constant, or capacitance (Supplemental Table 2). The similar membrane capacitance is consistent with the anatomical work, which shows that cells with the S403L mutation did not differ in soma size from WT *hTSC1* controls (Figure 2I). Analysis of spikes confirmed that cells classified as FS had electrophysiological properties typical of FS CINs in the neocortex, such as high-frequency discharge patterns with little spike-frequency adaptation (Average adaptation ratios: 0.61 for WT *TSC1* cells, 0.75 for S403L cells). Only one S403L cell was not capable of sustained AP discharge throughout a 1 second suprathreshold current injection. Interestingly, analysis of single APs evoked by a threshold current injection revealed a smaller fast afterhyperpolarization (fAHP) immediately following the AP in S403L than WT *hTSC1* cells (Figure 3C, 3D). Furthermore, we found that the peak medium AHP (mAHP) following a 1-sec spike train was smaller for S403L than WT *hTSC1* cells (Figure 3E, 3F). We found no differences in the functional properties between S403L and WT *hTSC1* cells classified as non-FS (Supplemental Table 2). Overall, the physiological data indicate that the S403L missense variant results in both a reduced fAHP and mAHPs in CINs classified as FS.

## Discussion

There is a growing number of missense variants in ASD. Herein, we built upon a successful *in vivo* screening assay (Vogt et al., 2014, 2015a, 2018; Hu et al., 2017a; Pla et al., 2018) to understand the impact of these variants in a group of cells implicated in ASD, CINs. We did this by examining a gene underlying a syndrome with high rates of ASD, i.e. *TSC1*, and that inhibits MTOR. We found that *TSC1* missense variants could complement a common phenotype used to assess *TSC1* dysfunction, i.e. increased cell size, suggesting that this measurement is a poor rheostat to assess what impacts these variants cause. However, a few variants were not able to complement the altered PV expression, suggesting that this measurement may be a better test of the effects induced by these variants.

Why two of the five variants were deficient at complementing the PV expression phenotypes is still unknown. However, they are found in mid to carboxy-terminal regions of Hamartin, which could imply distinct protein/protein interactions or unique functions of this region. Moreover, the S403L variant is a potential serine phosphorylation site that has not been reported in previous reports. Future studies will investigate whether this is a new kinase target and the impact that this mutation has on other cell types. In addition, these data support the idea that the central region and carboxy terminus of Hamartin might regulate processes that regulate PV expression, which will be tested in future studies

An interesting observation of this study is that the S403L missense *TSC1* variant resulted in a subtle reduction of the fAHP following a single AP and the mAHP following a train of APs in CINs with FS properties, with no apparent changes to passive membrane properties. This result differs from other *TSC1* studies, where cell-type-specific deletion of *Tsc1* in mice causes several electrophysiological alterations in both passive and active membrane properties (Normand et al., 2013; Kosillo et al., 2019; Malik et al., 2019). AHPs play an essential role in shaping neuronal firing properties (Hille 2001). Although we did not see any differences in the AP discharge properties of S403L compared to the WT *TSC1* FS CINs, as measured by spike half-width, spike adaptation, and the slope of the input-output curve, there could be other compensatory mechanisms at play. Our finding that expression of the S403L missense variant also increased the number of PV-expressing CINs could potentially relate to our physiological observation. PV is a calcium-binding protein that is found in specific classes of neurons, including CINs with FS properties (Celio, 1986; Wonders and Anderson, 2006) If MGE-derived CINs overexpress PV, it could modulate the intracellular calcium dynamics that occur during action potentials. Indeed, different calcium-activated potassium channels have been shown to be responsible for the various types of AHPs in mammalian neurons (Faber and Sah, 2003; Hille 2001).

Overall, these data demonstrate that unique variants in *TSC1* can impact the molecular and physiological properties of CINs. Moreover, it is intriguing that sensitive changes in distinct cell types can occur via single amino acid changes in *TSC1*, while more common phenotypes, i.e. increased cell size, is not observed. This is important in the context of our previous work, which showed that PV expression rapidly changed but cell size did not, when mouse *Tsc1* knockouts were treated with rapamycin and then allowed to recover (rapamycin removal) in a short time period (Malik et al., 2019). Thus, PV expression and CIN function may be sensitive readouts in TSC and a new way to understand the impact of the growing number of ASD variants. These data support the idea that mild changes in the function of critical cellular signaling events may be a factor influencing the development and mature functions of these unique neurons in TSC and potentially ASD.

## Ethics Statement

The animal study was reviewed and approved by the Michigan State University Institutional Animal Care and Use Committee.

## Author Contributions

DW performed primary cultures and analyses. SMB performed western blots. KA performed blinded cell counts. LEM performed physiology experiments. AMS and DV performed MGE transplants. DW, LEM, SRC and DV planned out experiments and wrote initial paper, all authors edited figures and text.

## Funding

R00-NS096108 (to SRC), R00-NS096108-S1 (to SRC and LEM). Spectrum Health-Michigan State Alliance Corporation (to DV).

## Acknowledgments

We would like to thank colleagues at Michigan State University and the Neuroscience Program for support and critiques.

## Notes

### Competing Interest Statement

The authors have declared no competing interest.

